# The mouse brain after foot-shock in 4D: temporal dynamics at a single-cell resolution

**DOI:** 10.1101/2021.05.03.442441

**Authors:** V. Bonapersona, H. Schuler, R.D. Damsteegt, Y. Adolfs, R.J. Pasterkamp, M.P. van den Heuvel, M. Joëls, R.A. Sarabdjitsingh

## Abstract

Acute stress leads to sequential activation of functional brain networks. The challenge is to get insight in whole brain activity at multiple scales, beyond the level of (networks of) nuclei. We developed a novel pre-processing and analytical pipeline to chart whole-brain immediate early genes’ expression – as proxy for cellular activity – after a single stressful foot-shock in 4 dimensions; that is, from functional networks up to 3D single-cell resolution, and over time. The pipeline is available as R-package. Most brain areas (96%) showed increased numbers of c-fos+ cells after foot-shock, yet hypothalamic areas stood out as being most active and prompt in their activation, followed by amygdalar, prefrontal, hippocampal and finally thalamic areas. At the cellular level, c-fos+ density clearly shifted over time across subareas, as illustrated for the basolateral amygdala. Moreover, some brain areas showed increased numbers of c-fos+ cells, while others dramatically increased c-fos intensity in just a subset of cells; this ‘strategy’ changed after foot-shock in half of the brain areas. All single-cell data can be visualized for each of the 90 brain areas examined through our interactive web-portal.

## Introduction

Acute stress leads to the activation of multiple functional brain networks, as demonstrated in humans using functional magnetic resonance imaging (fMRI, for reviews ^1,2^). Spatial resolution beyond the level of (networks of) nuclei, however, is currently not possible. This severely limits our understanding of stress-induced changes in activity at a whole brain level. For instance, are all cells within nuclei equally affected at various time-points after stress? Such a question cannot even be answered with high-field magnets in rodents.

The alternative to fMRI is the use of whole-brain microscopy which has excellent spatial resolution (~5μm, ^3^) yet very poor time resolution, usually confined to a single time-point. We here solved this conundrum by largely improving, expanding and changing a technique using immediate early genes (IEG) ^4^, in this case c-fos (Supplementary Note 1), which has been used as a post-mortem marker of cellular activity ^5^. Previous studies report that c-fos mRNA can peak at different times across brain areas after swim or restraint stress ^6^. This suggests that there may be multiple waves of c-fos activation throughout the brain, which could be used to map the temporal dynamics across all brain areas up to the level of single cells and from minutes to hours after the initial stimulation.

To understand how single cells across the brain in 3D adapt their activity at various time-points after stress, we exposed adult male mice to a single stressful foot-shock and charted cellular activity across the whole brain, using c-fos staining in 89 brain areas as a proxy thereof. To this end, a pipeline for data preprocessing and analysis at different spatial resolutions was developed, allowing investigation from the macro- (functional networks) to micro- (single cell resolution) scales and with a pseudo-time scale.

## Results

### Overview of the pipeline and quality control

Figure 1 summarizes the main features of the pipeline including experimental procedure, image processing (Step 1), data cleaning (Step 2), data pre-processing (Step 3) and analysis.

**Figure 1.**
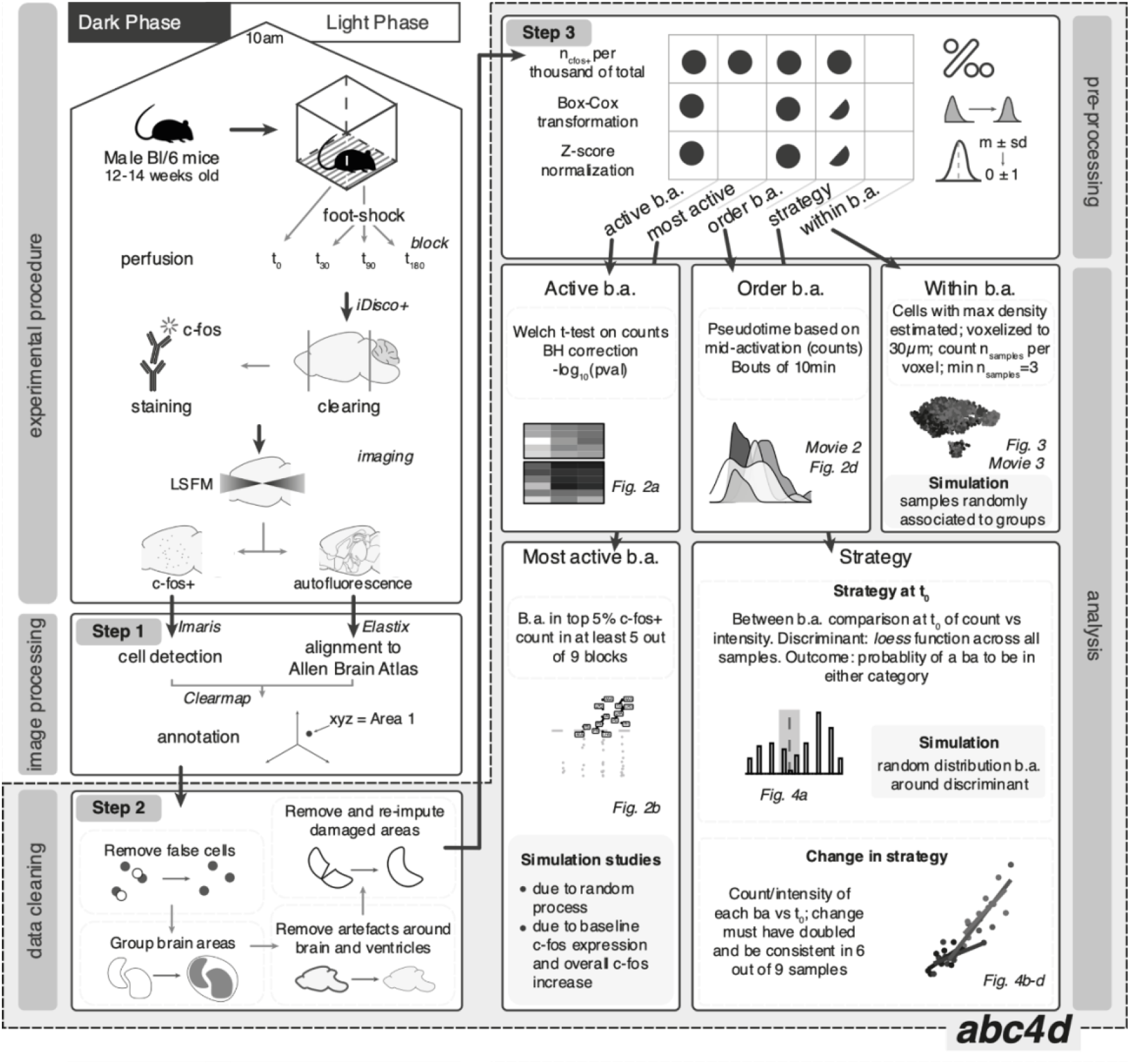
Schematic overview of the pipeline. Animals were perfused at different time-points (n_animals_ = n_time_ point*n_blocks_ = 4*9 = 36) after foot-shock. Whole-brain samples were processed with iDisco+ protocol; c-fos+ cells imaged with light-sheet fluorescent microscopy. Cells were detected with Imaris and annotated to the Allen Brain Atlas with Clearmap. Output yielded xyz coordinates per cell. Quality control (data cleaning) consisted of removal of various artefacts and of grouping brain areas (b.a.) to the spatial resolution of interest. Data preprocessing of b.a. was performed for each of the analyses. Circle: step required; half-circle: step recommended but not required. Strategy refers to t_0_ strategy categorization, as well as change of strategy over time. The frame at the bottom right summarizes the analyses conducted, and the main statistical decisions made. Data cleaning (Step 2), pre-processing (Step 3) steps as well as analyses are explained in detail in the Methods section and have been implemented in the *abc4d* package.

In brief, cell detection (Step 1) was performed with Imaris’s spot object (Supplementary Figure 1 a), after which it was aligned to the Allen Brain Reference Atlas (ABA) with Clearmap. Precision of alignment was assessed by comparing how sample images and template images would distort landmarks which were previously manually placed (Supplementary Figure 1 b). The average absolute difference was 8.39 ± 5.88 μm (mean ± SD) in the horizontal plane and 10.92 ± 12.44 μm (mean ± SD, maximal displacement = 23.36 μm) in the sagittal plane, with more laterally placed landmarks being less precise; i.e., on average an uncertainty of roughly one soma. Alignment did not differ per condition, suggesting that alignment error should not affect our results.

During data cleaning (Step 2), first unspecific binding was mitigated by removing background signals and applying a mask of 3 voxels (~75 μm) around the borders of the brain and ventricles (Movie 1), and by removing cells with abnormally high intensity (n_cells removed_ = 12). The background and mask step accounted for ~97% of the removed cells. Second, across all samples, ~5% of the brain areas showed some form of damage; these were removed from the analyses and re-imputed. Ultimately, the number of cells removed during the quality control procedure did not differ between groups (Supplementary Figure 2).

The next step in the pipeline is data pre-processing (Step 3). As summarized in the figure (Figure 1 upper left panel), data-preprocessing is specific for each analysis type. Of note, we used a block design, meaning that a ‘block’ (i.e., mice from the same cage, one animal for each time-point, processed simultaneously to avoid isolation stress) was the experimental unit of randomization and processing of samples. This type of design is essential for effective batch effect correction. The pipeline is available for similar future questions in the R package developed for the purpose, *abc4d* (“Analysis Brain Cellular activation in 4 Dimensions”), which is interoperable with several other annotation/alignment tools.

### Single cell activity is increased after foot-shock throughout the brain, but with spatial and temporal specificity

The total number of c-fos+ cells (n_cfos+_) across the brain was increased 30 minutes (t_30_) after foot-shock induction and remained elevated at t_90_ and t_180_. It returned to to levels 300 minutes after foot-shock (Supplementary Figure 3). Across batches, n_cfos+_ was comparable to that of previous literature ^3,7^. Of note, t_0_ animals were placed in a foot-shock chamber but did not receive a foot-shock. As a consequence, they should be considered as a “mildly stressed” (novelty stressor) group rather than true baseline controls.

To test the extent of c-fos+ expression throughout the brain, we performed pairwise comparisons (Welch t-test, one sided) between each foot-shock time point (t_30_, t_90_, t_180_) and t_0_ (Figure 2A). 86 out of the 89 brain areas had a significant increase in c-fos+ cells in at least one of the time points. Only three brain areas were not significantly changed, i.e. medial preoptic nucleus, ventral anterior-lateral complex of the thalamus, and ventro-posterior complex of the thalamus. The time point t_180_ had the highest number of significant brain areas (n_sig brain areas_ = 85), followed by t_90_ (n_sig brain areas_ = 79) and t_30_ (n_sig brain areas_ = 40). The effect sizes (*g*±SD) ranged between −0.32±0.23 (Midbrain raphe nuclei, t_30_ vs t_0_) and 5.17±0.96 (Subiculum, t_90_ vs t_0_) with a mean of 1.62±0.57.

**Figure 2.**
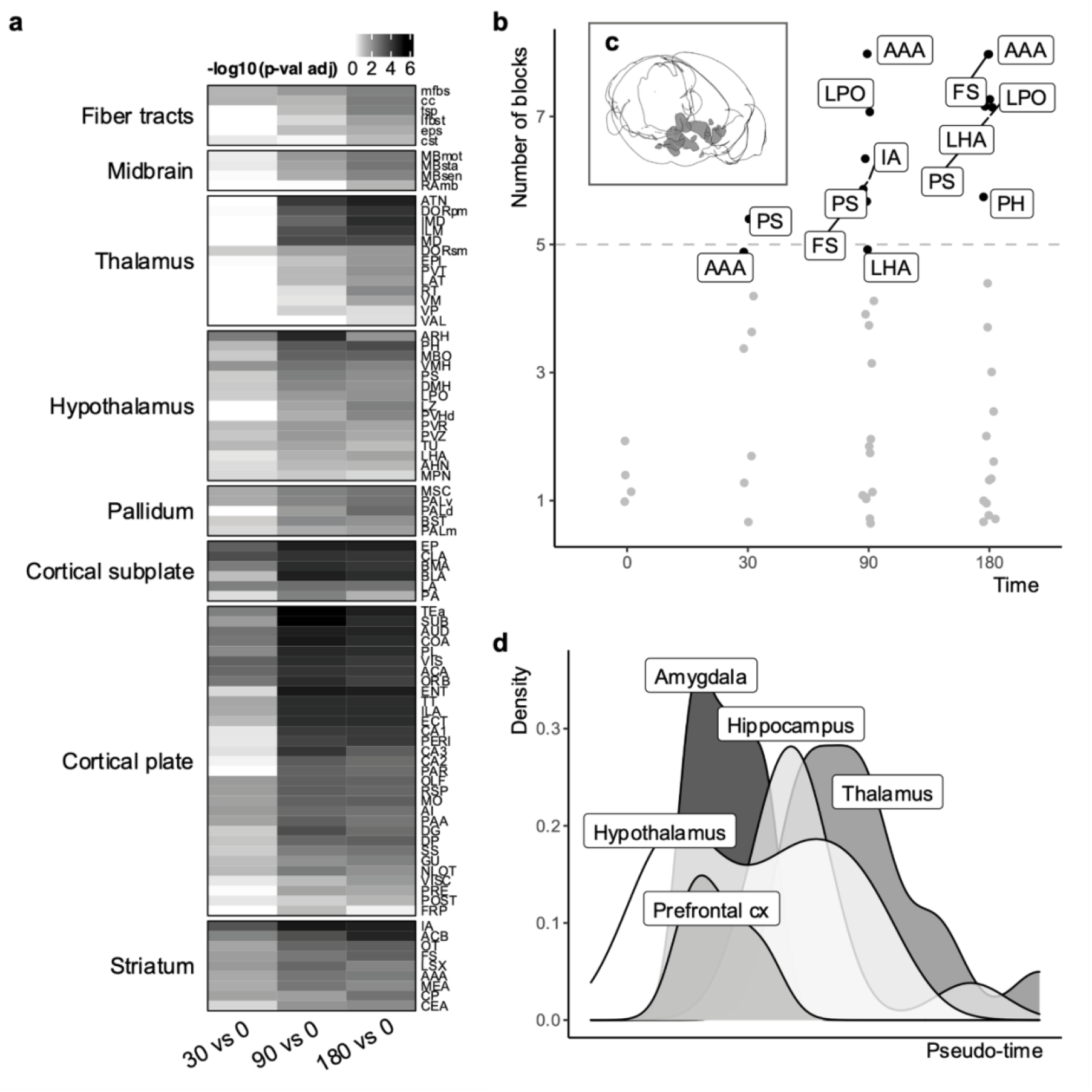
Most brain areas are activated by foot-shock, with spatial and temporal specificity. a) Heatmap of -log_10_ p-values derived from pairwise comparisons of each foot-shock time point (t_30_, t_90_, t_180_) against t_0_ for each brain area. White: p-val_adj_ >=0.05; grey_shades_: p-val_adj_ < 0.05. The legend numbers correspond to the log values. b) A set of hypothalamic areas was consistently found to have the highest (top 5% of the distribution) number of c-fos+ cells (per thousand of total). The criterion for consistency was 5 out of 9 samples. Abbreviations are explained in Suppl. Table 2. c) Cartoon of the brain areas identified in b; created with *brainrender* ^8^ d). Functional order of brain areas’ c-fos activity following foot-shock. Brain areas were ordered based on a pseudo-time depending on c-fos+ activation across the time points, and grouped based on functional categorization important for the stress response ^9^. Hypothalamic areas are the first to reach the mid-point of their activation, followed by amygdalar, prefrontal, hippocampal and lastly thalamic areas. For an interactive visualization of the single brain areas rather than the categorization, see Movie 2.

Since nearly all brain areas were active in at least one time point, we aimed to identify which brain areas were more active than others. To answer this question, we identified for each block (i.e., a unique set of each time-point) the brain areas that had the highest (i.e., top 5% of the distribution) c-fos+ cell count density (per thousand of total, n_cfos+/tot_). Under random circumstances, the same brain area would be in the top 5% in at least 5 out of 9 samples in about 1% of the cases, as illustrated by a simulation study (Methods). Being selected by at least 5 samples was therefore used as a criterion to define consistency of highly active brain areas. In our experimental data, the criterion was met by 8 brain areas (Figure 2 b), which belonged mostly to the hypothalamus (Figure 2 c). This number (n_highly active_ = 8) was much higher than the 1% expected (n_randomly active_ = 1) by sheer randomness. With a simulation study, we confirmed that this activation could also not be attributed to the spatial localization of c-fos throughout the brain, as reported by the Allen Brain Atlas (Supplementary Figure 4).

Next, we hypothesized that although in most brain areas the number of c-fos+ cells is increased after foot shock, the peak of activation would not occur at the same time for every (network of) brain area(s). Brain areas are expected to be involved at different stages – and therefore at different times – of the stress response as earlier proposed, based on human fMRI studies ^1^. We organized brain areas on a pseudo-time scale, based on the time-point in which a brain area (median across blocks) would reach the middle of its activation. This pseudo-time should only be interpreted relatively. We visualized the order of activation (Figure 2 d) of (networks of) the brain areas, using a functional categorization valuable to the stress response ^9^. Based on this classification, hypothalamic areas were found to be activated first, followed by amygdalar and prefrontal, hippocampal, and finally thalamic areas. Movie 2 shows a visualization of all brain areas over time.

### Time-dependent wave of activation within the Basolateral Amygdala

While the results so far confirm – in rodents – insights at the network level earlier obtained in humans with fMRI ^1^, our main goal was to investigate dynamic brain activity after foot-shock with higher spatial resolution, up to the single cell level, which is a great advantage over e.g. fMRI studies. Rather than highlighting all areas, we here illustrate the findings for the basolateral amygdala (BLA), an area key for the cognitive processing of a foot-shock ^10^ and stressful conditions in general ^11^; for the remaining areas we refer to an open-source user interface (https://utrecht-university.shinyapps.io/brain_after_footshock/), with which one can browse all other regions.

We hypothesized that the increase of c-fos+ cells was not uniform across the BLA; but, rather, may be restricted to different sub-parts or cells. For each sample independently, we identified the most densely activated part of the BLA, i.e., the part with the highest number (density) of c-fos+ cells relative to the rest of the BLA. All samples considered, there are obvious regional distributions across time points (Figure 3 a, Movie 3), which are not evident when samples are randomly associated to the experimental groups (Supplementary Figure 5 a).

**Figure 3.**
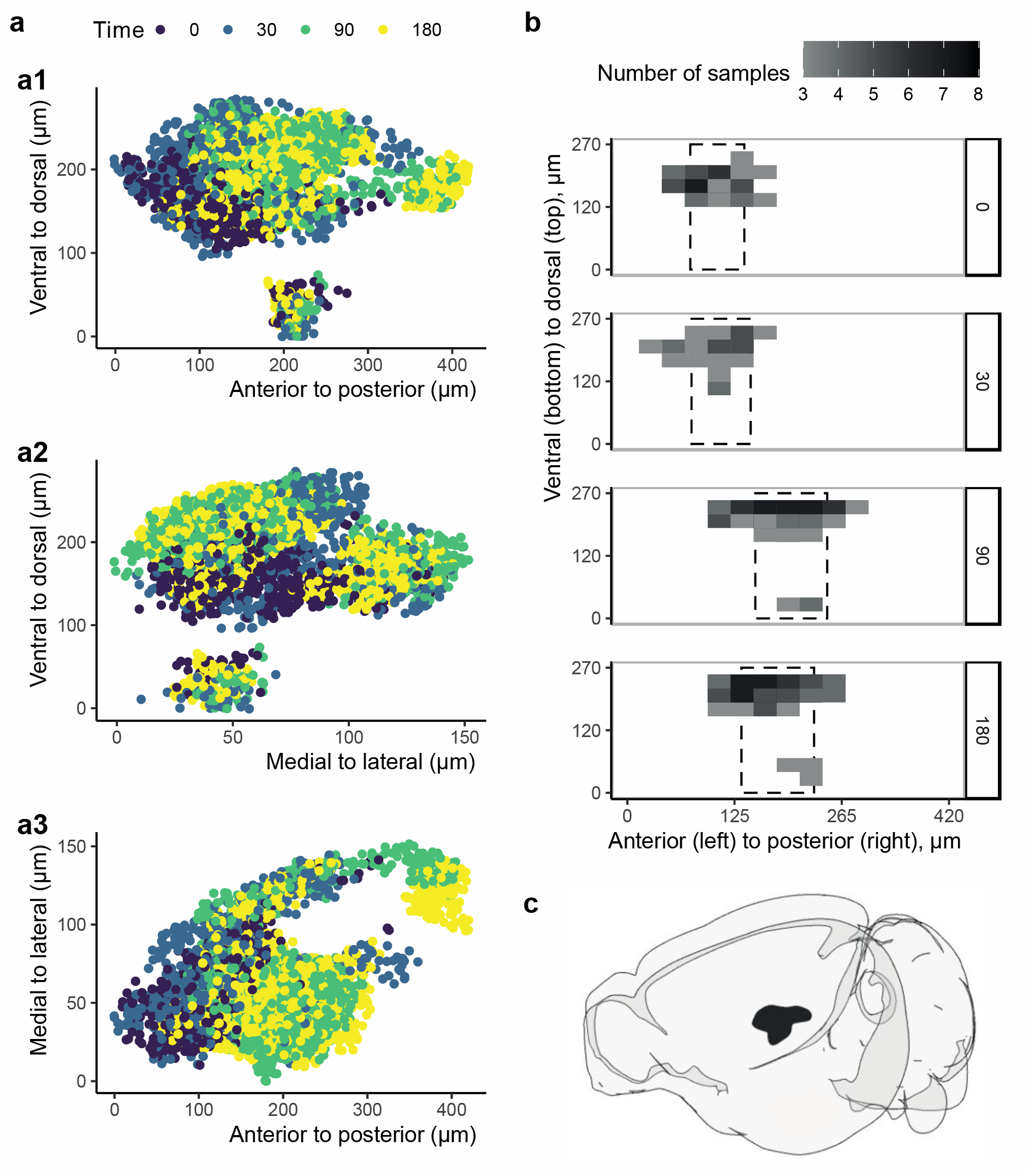
Changes in c-fos+ cell density within the Basolateral Amygdala. a) Cells in high density regions of the right BLA. The 3D cell coordinates are represented as a set of three 2D graphs, one for each couple of coordinates (xy, yz, xz). Each dot is a cell of a sample in a region with highest density. The colors refer to the different time points. b) The densest c-fos+ sub-part of the BLA moved from more anterior (t_0_ and t_30_) to more posterior after foot-shock (t_90_ and t_180_). The BLA has been voxelized (voxel size 30×30×30μm), and the fill color refers to the number of samples with at least one cell in that voxel. The dashed box indicates the mean and SD per group along the posterior-anterior axis. c) Cartoon visualization of the right BLA in the same orientation of a1 and b, created with *brainrender*^8^.

We voxelized the xyz coordinates (voxel size: approx. 30μm x 30μm x 30μm) and visualized per time point which voxels have at least one cell from three different samples. As shown in Figure 3 b, at later time-points after foot-shock, i.e. t_90_ and t_180_, the highest density of c-fos+ cells in the BLA was found to be more posterior (difference of ~96μm from anterior to posterior, 23% of BLA) than at t_0_ and t_30_.

### Cells use different strategies of activation, which can change after foot-shock

With 3D microscopy, one can not only count the n_cfos+_, but also quantify the intensity of c-fos staining per cell. Among other parameters, a cell is considered c-fos+ if the intensity of c-fos is higher than the background (i.e., signal-to-noise ratio). As a consequence, one would expect the n_cfos+_ per brain area to strongly correlate with the average c-fos intensity. In other words, brain areas are expected to be normally distributed along the correlation line, as shown by our simulation (Supplementary Figure 5 b).

However, this was not the case (Supplementary Figure 5 c). Rather, different brain areas have a preferential strategy of activation: a few very active (i.e. low count, high intensity) cells, versus many lowly active cells (i.e. low intensity, high count). We categorized the brain areas of each t_0_ sample (corresponding to a very mildly stressful condition) based on their ‘strategy’, i.e. their preference for increasing in count or intensity. Across samples, we then calculated the probability of each brain area to belong to either categorization (Figure 4 a). The results showed a bimodal distribution (Figure 4 a), which is clearly different from the normal distribution expected under our hypothesis (Supplementary Figure 5 b). Therefore, whether a brain area is activated by increasing n_cfos+_ or by increasing the average intensity of c-fos per cell is unlikely to be the result of a technical characteristics or of a random process.

**Figure 4.**
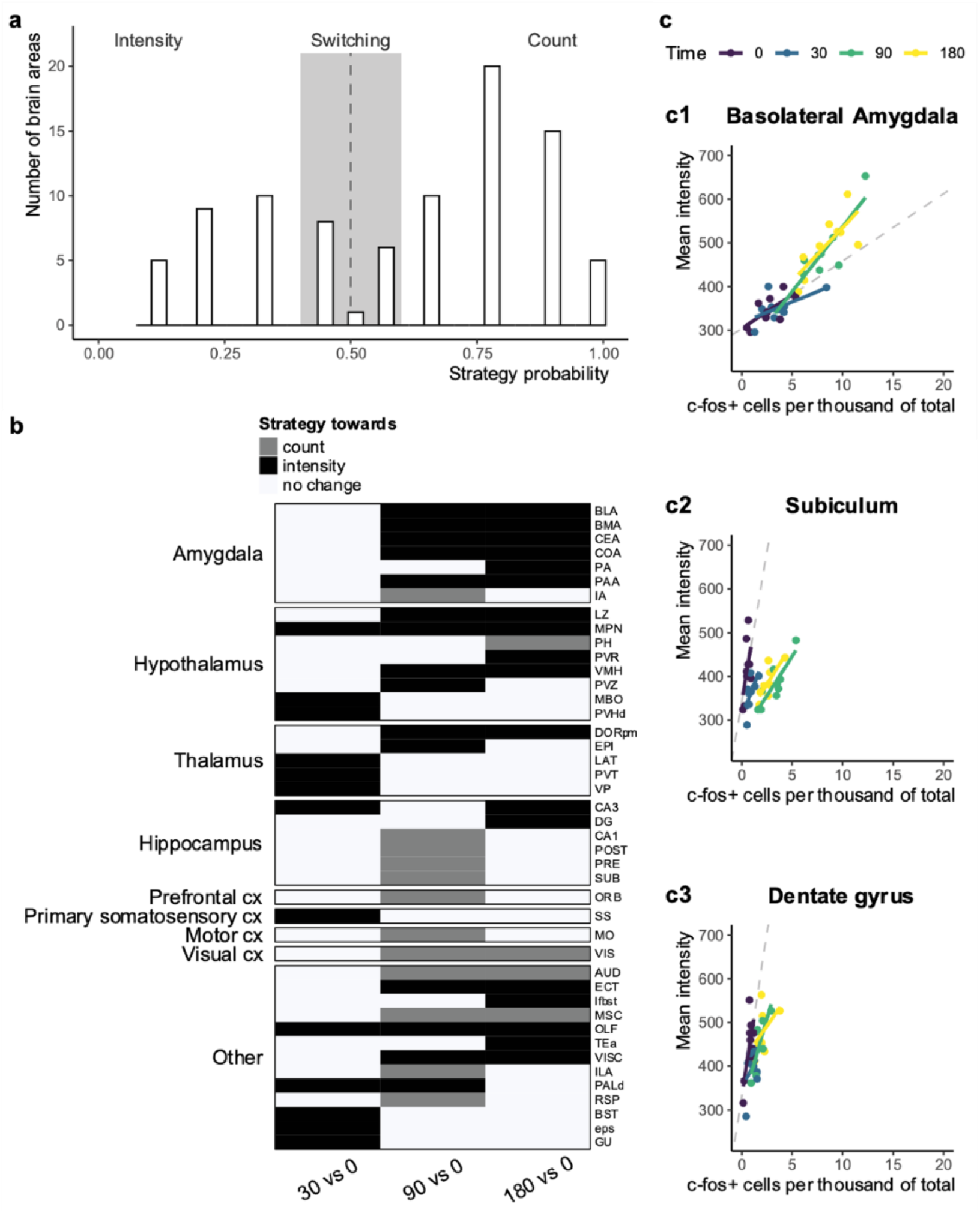
Brain areas have a preferential strategy of activation, which may change after footshock. a) Preferential strategy of brain areas based on t_0_ data, i.e. the relationship between intensity and count per brain area. Histogram of the number of brain areas across the strategy probability. If a brain area would be activated by indiscriminately increasing c-fos+ cells and expression, the distribution would be Normal (Supplementary Figure 5b), with μ around 0.5. The bimodal distribution suggests that certain brain areas preferentially increase the number of c-fos+ cells (probability > 0.5, count), whereas other increase the mean c-fos expression (probability < 0.5, intensity). b) The strategy of brain areas can change after foot-shock. Binary heatmap of how strategy can change across brain areas for pairwise comparisons of time points. White corresponds to no change in strategy, black to a change towards intensity and grey to a change towards count. Of note, a change towards intensity does not necessarily mean that the brain area does not increase count; rather, it means that the increase in intensity cannot be explained by the increase in count alone. c) Representative examples of a brain area that after foot-shock changes strategy towards intensity (BLA, c1) or count (Subiculum, c2). The dentate gyrus (c3) was added for literature validation (see Discussion). Each dot represents one sample; the line represents that correlation between count and intensity per group. Dashed line represents what one would expect based on t_0_ activation.

Intensity and count are therefore expected to be related within brain areas, rather than across the whole brain. This relationship should be constant across all groups; if not, foot-shock must have induced transcriptional changes in specific subsets of cells.

We therefore examined whether the strategy of a brain area changes after foot-shock, relatively to t_0_. For each time-point after foot-shock (t_30_, t_90_, t_180_), we selected brain areas with a consistent (at least 6 out of 9 samples) change in either count and intensity, and calculated to what extent count and intensity were increasing compared to each other. We categorized brain areas as “changing strategy” if they at least doubled the increase in one category. 43 brain areas met these criteria (Figure 4b). Of these, 30 increased activation by means of intensity rather than c-fos+ cell count, especially in the amygdala, hypothalamus and thalamus. Of note, the increase in intensity for the amygdalar nuclei was present only for the time points t_90_ and t_180_. Figure 4 c displays two brain areas (BLA and Subiculum) as representative examples of activation strategy towards intensity and count, respectively. We also added a visualization of the dentate gyrus (Figure 4 c, part 3) as a validation, since this area has been described to increase intensity of IEG staining after stress in a very limited subset of cells ^12,13^.

## Discussion

Human fMRI studies over the past decades have shown that acute stress activates multiple functional brain networks, with hypothalamic and amygdalar areas being among the first to be activated, and areas linked to higher cognitive functions such as prefrontal cortex and hippocampus following in due course ^1^. Yet, beyond the level of (networks of) nuclei, e.g. up to single cells, very little is known about stress-induced effects at a whole brain level. Are all cells among or within nuclei equally affected at various time-points after stress?

To address this question, we first had to develop a thorough and robust analytical pipeline, investigating changes in cellular activity after a highly stressful foot-shock in mice over time, by staining for the IEG c-fos and introducing a pseudo-time metric; this is summarized in Figure 1. Several tools had been developed earlier to transform images into machine-readable data (for example, ^3,14,15^), but no publication so far has thoroughly explored the subsequent steps of data analysis, i.e. dealing with missing values, batch effect corrections, normalization and transformation. Also, so far analyses were limited to a region-based approach, where the number of active cells is calculated for each brain area separately. Voxel-based analyses similar to MRI have only recently been developed ^16^. Here, we took this one step further, and suggest analyses for *i)* the time dynamics and *ii)* the single cell level. The resolution up to the level of single cells is certainly one of the major advantages of whole-brain microscopy. In the future, c-fos+ cells could be characterized in more detail, being able to distinguish excitatory from inhibitory neurons, or neurons from glia cells, as the current findings confirm that c-fos staining is not confined to neurons only (e.g. ^17^). This could be achieved by multiple concurrent stainings, or by computationally categorizing cellular morphology ^18^. Our pipeline can be applied independently of the type of software used for alignment and annotation, and it can analyze 3D (i.e. whole-brain) or 4D (over time) experimental designs. Furthermore, it is interoperable with several other annotation/alignment tools. It is available in the newly developed R package *abc4d*, to which new improvements can be easily added in future. *Abc4d* also includes a framework of simulation studies, where the null hypotheses for different analyses can be investigated. An overview of the package is provided in the cheat-sheet (Supplementary Figure 6).

With this toolbox in place, we addressed our main question. Although foot-shock increased the activation of 96% of the brain areas, distinct temporal dynamics in networks of brain areas stood out. Thus, foot-shock first activated (cells in) hypothalamic areas, followed by amygdalar and prefrontal, hippocampal and lastly thalamic areas. This is largely in line with the earlier human literature using fMRI ^1^. Additionally, we demonstrated within the basolateral amygdala – a key area in the processing of a stressful foot-shock ^10^,^11^ –, a clear shift after foot-shock from activation of cells in the lateral-anterior part towards a more posterior-medial subset of cells. Although we here present information about the basolateral amygdala only, information about all other areas investigated is available for closer scrutiny. To dive deeper into any area of interest, we provide an interactive interface on the data at our web portal (https://utrecht-university.shinyapps.io/brain_after_footshock/). Lastly, our approach allowing single cell investigation revealed that brain areas follow specific c-fos expression strategies that are skewed towards either an increase in number of c-fos+ cells or c-fos intensity per cell. Importantly, in a subset of areas the strategy changed after foot-shock.

The finding that brain areas use a distinct strategy in c-fos cell activation under mildly stressful conditions such as a novel environment (in this case the shock-box at t_0_) compared to exposure to a very stressful situation like an inescapable foot-shock is very novel. One can currently only speculate about the functional relevance of the two main cellular ‘strategies’. Earlier studies showed that the expression of c-fos is proportional to the rate of firing of the cell ^19^. If so, one could hypothesize that in certain brain areas many cells are slightly more activated after stress (i.e. express c-fos above detection threshold), whereas other brain areas may have only a few cells that are very strongly excited, which would lead to an increase in their c-fos intensity. Of note, in amygdalar areas the increase in intensity was particularly observed at 90 and 180 minutes after foot-shock, which is compatible with a gene-mediated, possibly glucocorticoid-dependent mechanism ^20^. This observation that a limited number of cells gets highly activated fits extremely well with current views on engrams ^21^. Using inducible IEG promotor approaches in areas of interest, others have shown that immediately after contextual fear conditioning – also involving foot-shock exposure – a consistent ~10% of (baso)lateral amygdala neurons becomes part of the engram, while participation of dentate granule cells is much lower ^22^. This resembles the ~5% of BLA neurons with very high intensity staining we observed 90-180 minutes post foot-shock and the far lower number of high-intensity c-fos+ cells in the dentate. The somewhat lower percentage of c-fos+ cells in the BLA we observed may be explained by the recency of the foot-shock ^22^, the level of excitability prior to foot-shock ^23^ and/or the fact that we used c-fos rather than arc as IEG. Very high intensity of IEG staining in a small subgroup of dentate cells was also reported after swim stress, which was found to increase DNA demethylation in the dentate gyrus at specific CpG sites close to the c-fos transcriptional start site, in the gene promoter region of early growth response protein 1 ^12,13^.

There are some limitations to consider. The choice of c-fos as an activity marker is arguably appropriate in the case of acute stress exposure ^5^, but it is by no means the only IEG one could choose for the current approach. Other markers of cell activity may afford additional insights into the circuits being activated after stress. For example, *Arc* and *Egr1* were reported to be transcriptionally activated following acute stress in a multi-omics approach ^24^. Ideally, a combination of markers should be applied to get a completer view. Another technical limitation is linked to the current size restraints of imaging with light-sheet microscopy. In our study, we trimmed the most frontal and most caudal parts of the brains. Although this does not impact the current methodology and the main finding of between- and within-area cellular differentiation in response to stress, some brain areas involved in the acute stress response (e.g. locus coeruleus) are missing. A solution could be to divide the brain for scanning, but to analyze the data together, after the appropriate corrections. A third consideration concerns the pseudo-time approach. On the one hand, it overcomes the absence of high frequency sampling (as possible with fMRI). On the other hand, it is a mere approximation of real-time processes. Interestingly, at the macro-scale, the order in which brain regions are activated following the acute stress of a foot-shock strongly resembles that seen with fMRI after acute stress in the human brain ^1^. This lends credibility to our approach and also emphasizes its translational potential. Importantly though, while it might be an acceptable approximation of the activation phase, the method gives little insight in the gradual turning off of brain areas, since this also depends on the half-life time of the c-fos protein. The half-life time may differ across brain areas ^5^, something that could be investigated with a meta-analytic approach.

Despite these limitations, the ready-to-use pipeline for 4D immunohistochemical whole-brain analysis presented in this report (and supported by a new R package) revealed that stressors like an acute foot-shock not only sequentially activate functional networks in the brain, but also specifically activate subsets of neurons, using different strategies of activation.

## Acknowledgments

We would like to acknowledge the MIND facility (UMC Utrecht Brain Center) for 3D imaging by lightsheet microscopy (mindresearchfacility.nl), and Roger Koot for the IT infrastructure. We would like to thank Kevin Kemna for suggesting the creation of the R package and the helpful feedback on the analyses. This work was supported by the Consortium of Individual Development (CID) and BRAINSCAPES, which are funded through the Gravitation program of the Dutch Ministry of Education, Culture, and Science and the Netherlands Organization for Scientific Research (NWO grant numbers 024.001.003 and 024.004.012).

## Author contributions

V.B. contributed with the conceptualization, data curation, analysis, investigation, methodology, software, visualization and writing the manuscript; H.S. contributed with the methodology, performance of experiments, analysis and reviewing the manuscript; R.D. contributed with the performance of experiments; Y.A. contributed with support with experiments and reviewing the manuscript; R.J.P. provided infrastructure, contributed with funding acquisition and reviewing the manuscript; M.P.vdH. contributed with the conceptualization of the project and reviewing the manuscript; M.J. contributed with the conceptualization, funding acquisition, project administration, supervision, and writing the manuscript; R.A.S. contributed with the conceptualization, project administration, supervision and writing the manuscript.

## Methods

### Animals and husbandry

8 to 10 weeks old male C57Bl/6JOlaHsd mice were purchased from Envigo (Harlan, France) in 3 separate batches. The animals were habituated to the animal facility for at least two weeks, then tested at the age of 10 to 14 weeks. Until the experimental day, the animals were housed in groups of five in type II L cages (365×207×140mm, Tecniplast®) on a 12h dark/light cycle (light phase: 9.00AM–9.00PM), 22±2 °C, humidity at ± 64%, with *ad libitum* access to standard chow (Special Diet Services, UK, sdsdiets.com) and tap water. Experimental cages were placed on racks without a specific order and left undisturbed in the same experimental room, except for weekly cleaning by animal caretakers unfamiliar with the study design. A copy of the work-protocol (license: 527/16/644/01/06, 527/18/4806/01/01) as well as a step-by-step protocol can be found on the Open Science Framework page of the project (osf.io/8muvw). All animal procedures were approved by the Animal Ethical Committee at Utrecht University (license: AVD1150020184806), the Netherlands. Every effort was taken to minimize animal suffering in accordance with the FELASA guidelines and the Dutch regulation for housing and care of laboratory animals.

### Experimental design

The primary aim of this study was to develop a methodology to be able to analyze brain-wide activation over time, specifically of the stress system. The experiments were designed to address batch effects, missing values and normalization. An overview of the experimental procedure, data cleaning, preprocessing, and analysis can be found in Figure 1.

The main experiment used a uniform block design, where each block corresponded to a separate cage of animals. Each cage contained an animal of each experimental group (being ‘time after foot-shock’ and control; n_time point_ = 4). The animals were killed at three different time points after receiving a foot-shock (n_time point foot-shock_ = 3). A control group underwent the same identical procedure but did not receive the foot-shock. This group is considered time point 0 min (t_0_). Of note, this control group did experience a novel environment, and its activation should therefore be considered not as ‘baseline’ but as mildly stressed. The time points refer to the moment of transcardial perfusion, and were chosen to comprehensively model all phases of the stress response. They cover the initiation (t = 30 min, t_30_), maintenance (t = 90 min, t_90_) and termination (t = 180 min, t_180_) of the HPA axis response, while considering the required time to synthesize c-fos ^5^ as well as the delay of 30 minutes of increase of corticosteroid in brain tissue compared to blood ^25^. Randomization of the experimental groups occurred within the cage, and samples of the same block were processed together both pre- and post-mortem. In other words, each experimental group was represented in every four animals (block), but the order of experimental groups differed across blocks. This type of design is important for batch effects’ correction.

The following control experiments were performed. A 15 min time point (n_animals_ = 3) to validate that indeed 30 min is the earliest time point to detect an increase in c-fos protein expression. A 300 min time point (n_animals_ = 2) to validate that c-fos expression eventually decreases. A no primary antibody staining (n_animals_ = 3) to estimate the extent of unspecific secondary antibody binding. A home-cage control group (n_animals_ = 3) was used to validate that the c-fos expression of the t_0_ group was due to the experimental procedure (i.e., placement into a novel cage). To minimize the sample size of animals used, the animals of these control experiments were processed together with those of the main experiments, so that the comparison groups could be reused. As a consequence, these control experiments are only qualitatively assessed, and no formal statistical analysis is performed to avoid unnecessary multiple testing. Data of the control experiments is presented paired to the t_0_ group of the same batch.

Initially, we aimed to include males as well as females (see work protocol at osf.io/8muvw) in the experiment. However, the percentage of animals lost due to unforeseen circumstances (due to the antibody) was much larger than anticipated (30% instead of 10%). That is, the secondary antibody was less stable than expected; as a consequence, the quality of the scans was often insufficient for a brain-wide quantification. We therefore simplified the experimental design and used only 9 male mice per experimental group (n_block_ = 9, n_time point_ = 4, n_animals_ = 36). These were processed in three different batches of 3, 4, and 2 blocks, respectively. We chose males instead of females because more male samples had sufficient quality after the first two batches. This qualitative assessment took place before c-fos quantification. The sample size of 9 animals per group was in line with or larger than manuscripts previously published using whole-brain microscopy ^3,26,27^ Furthermore, it was sufficiently large to identify, study and mitigate batch effects. No experimental animal was excluded from the study for reasons other than insufficient staining quality. For a full description of the missing animals, see Supplementary Table 1.

### Foot-shock induction

At any given experimental day, a cage was brought to the experimental room. Each animal of the cage was placed in a separate foot-shock box (floor area: 250×300mm) at the same time. Since the animals were not earmarked, selection bias was limited by randomizing the order of the shock boxes when placing the animals. In this way, we aimed to limit a “shock box” specific effect. After 60 seconds, the experimental animals received a single foot-shock (0.8 mA, 2 sec). The t_0_ animal of each block was always placed in a “sham shock box”. This box was identical to the others, but it did not give a shock. After another 30 seconds, all animals were removed from the shock boxes. The t_0_ animals were euthanized immediately, whereas animals of the foot-shock groups were single-housed in new cages waiting for transcardial perfusion according to their specified time point. The new cages were enriched with *ad libitum* chow and water, as well as bedding material of the home cage to limit arousal due to a novel environment.

To acquire meaningful, yet above threshold c-fos expression, the optimal t_0_ condition (home-cage vs novel environment) and foot-shock intensity (0.4 mA vs 0.8 mA, assessed at t = 90 min) were established in a pilot study. The results of the pilot study were only qualitatively assessed.

The investigator (RD) performing the foot-shock procedure and the perfusion was not blinded to condition, since she needed to confirm that the foot-shock was successfully applied and that animals were perfused at the correct time point.

### Perfusion and tissue preparation

Euthanesia was performed with an intraperitoneal injection of 0.1 mL pentobarbital (Euthanimal 20%, 200 mg/mL) ~10 minutes prior to transcardial perfusion. The animals were perfused with ice-cold 1x PBS until blood clearance, followed by perfusion with ice-cold 4% PFA/1x PBS. Brains were extracted from the skull of perfused animals and stored in 4% PFA/1x PBS overnight for post-fixation.

Brains were cleared and stained following the iDisco+ protocol ^3^. Briefly, 24h post-fixation, samples were washed with 1x PBS. The olfactory bulbs and cerebellum were trimmed. A methanol/H_2_O gradient was applied to dehydrate the tissue, followed by decolorization with 5% hydrogen-peroxide. After rehydration with a methanol/H_2_O gradient, brains were permeabilized and remained in a blocking buffer for two days. Samples were then incubated in the primary antibody for seven days (rabbit anti c-fos, Synaptic Systems, cat. 226003, 1/1000 in 92% PTwH / 3% Donkey Serum / 5% DMSO). Following washing with 1x PBS / 0.2%Tween-20 / 0.1%Heparin (PTwH), brains were incubated for another seven days in the secondary antibody (Donkey anti-Rabbit IgG (H+L) Alexa 647™, Thermo Fisher Scientific, cat. A31573, 1/1000 in 97% PTwH / 3% Donkey Serum). Lastly, samples were washed with PTwH, dehydrated with a methanol/H_2_O gradient, incubated in 66% DCM / 33% Methanol, washed with 100% DCM, and then cleared and stored in the clearing agent dibenzyl ether (DBE), protected from the light. The total time required to complete the protocol is 25 days. The investigators processing the tissue samples were blind to the experimental groups.

### Imaging with Light-Sheet microscopy

Starting three days after clearing, samples were imaged with a light-sheet microscope (UltraMicroscopeII, LaVision BioTec), equipped with an MVPLAPO 2x/0.5NA objective (Olympus), an MVX-10 Zoom Body (Olympus) and a 10mm working distance dipping cap. The images were recorded with an sCMOS camera (Neo 5.5 sCMOS, Andor Technology Ltd; image size: 2560×2160 pixels; pixel size: 6.5×6.5μm^2^) using the software ImspectorPro (v.5.0285.0; LaVision BioTec). The samples were scanned in horizontal slices (step size: 3μm; effective magnification: zoom_body objective_ + dipping lens 2×0.63×=1.26×; sheet width: 60%) with two-sided illumination using the built-in blending algorithm. Two image stacks per sample were taken consecutively, without moving the sample in between recordings. This is essential for the later correct annotation of c-fos positive (c-fos+) cells to brain areas. To record the fluorescence of c-fos+ cells, we used a Coherent OBIS 647-120 LX laser (emission filter: 676/29). The images were recorded at 70% laser power, sheet NA of 0.076 (results in a 10μm thick sheet), and an exposure time of 100 msec, as well as with horizontal focus to reduce z-slice distortion (steps: 20). To highlight the intrinsic fluorescence of the tissue for registration of the sample to a template, we imaged with a Coherent OBIS 488-50 LX Laser (filter: 525/50nm). The images were recorded at 80% laser power, sheet NA at 0.109 (results in a 7μm thick sheet), and exposure time of 100 msec, without horizontal focus.

The investigators (VB, HS) conducting imaging and image processing were blind to condition. Samples were imaged in ascending order, with sample numbers being randomly assigned during perfusion.

### Image processing: cell detection

c-fos+ cells were detected from the 647nm image stack using Imaris (v9.2.0, Bitplane). A spots object was created and parameterized to detect cells with estimated xy-diameter of 10μm and an estimated z-diameter of 35μm (to avoid overcounting cells due to z-plane distortion). Thereafter, detected cells were filtered by quality (>18) and required to cross a threshold of minimum intensity (>225). The quality filter verifies that the shape of the detected cells aligns to the spots object within a threshold specified automatically by the algorithm. These parameters were optimized on pilot imagined brains to achieve optimal signal-to-noise ratio while avoiding ceiling effects of the cell intensity parameter. The xyz-position of each cell was exported for later annotation to a reference atlas.

### Image processing: alignment and annotation

Each cleared brain was registered to the Allen Mouse Brain 25 μm reference Atlas ^28^ by using ClearMap ^3^ as an interface to the open-source software Elastix v5.0 ^29^. Registration was performed using the autofluorescent images (488nm image stack). As per our setting, the images were first rotated, sheared and scaled with affline transformation, then translated onto the reference atlas with b-spline transformation.

The transformation matrix as calculated by Elastix was applied to the xyz coordinates of the Imaris detected cells using Transformix. In this way, detected cells were transformed into a template space, and each c-fos+ cell was assigned to a brain region. This was possible because each sample remained in the same position for both image stacks.

To estimate the error in the approach, we drew in ImageJ (Fiji) artificial spots (n_spots_ = 72) in both hemispheres (n_animals_ = 12) in three different locations (i.e., within dentate gyrus, mammillothalamic tract, amygdalar capsule). To estimate the error of the alignment, we calculated the distance between the expected position of the artificial spots and the position resulting from the alignment procedure.

The output for further data analysis is the xyz-position of every cell with their corresponding area code. The code was then translated to the respective brain area according to the brain region table provided by Renier and colleagues ^3^, which follows the hierarchical organization of the Allen Brain Atlas (ABA).

At this point, images have been transformed in machine-readable numbers. Other tools can be used until here. To continue with the following steps, one is only required to have files with xyz coordinates for each cell, together with their annotated brain area.

### Quality control and data pre-processing

Quality control, data pre-processing and analysis were planned on a subset of the data (n_blocks_ = 3), and then later extended to the full dataset. The experimenter coding the analyses (VB) was blinded to the experimental condition.

To mitigate technical noise, a series of quality control steps were performed on the xyz annotated coordinates of c-fos+ cells. We removed (false positive) cells from brain areas in which no counts were expected, either because these areas contain no brain tissue (background, ventricular system) or because they were trimmed from the sample (olfactory bulb, cerebellum, hindbrain). Next, we grouped the highest resolution areas of the ABA in line with the ABA hierarchical organization. The aim was to preserve as much spatial specificity as allowed by alignment inaccuracies in areas likely to be stress or c-fos sensitive, while minimizing the total number of brain areas to ease interpretation and to avoid unnecessary subsequent multiple testing. Accordingly, the categorization considered the region-specific distribution of glucocorticoid receptors as well as the region-specific c-fos expression after acute stress ^6,30)^. The hierarchical relationship of ABA areas is not complete, meaning not all larger brain areas can be fully subdivided into smaller brain areas. These “left over” spaces were removed from the analysis since they were deemed not interpretable.

An illumination artifact was present in all samples on the outside borders of the brain and the ventricles, presumably due to unspecific antibody binding. Initially, we aimed to use the no primary antibody control group to correct for this artifact; however, this was not possible as the number of c-fos+ cells in the no primary antibody control group was minimal. As an alternative solution, a mask of 75μm thickness was modeled along the inside border of the brain and the ventricles of the aligned samples (Supplementary Figure 3 a), and cells that fell within the mask coordinates were removed from further analysis. The size of the mask (25 through 175μm) was piloted in 3 samples. Ultimately, 89 brain areas were included in the analysis (Supplementary Table 2).

Lastly, we removed xyz coordinates with extremely high c-fos intensity. We qualitatively assessed histograms of the maximum intensity of c-fos+ xyz coordinates per brain area, and compared them across samples to identify potential unspecific binding of the protein (“spots”). The potential candidates had 2- to 10-fold higher intensity than others within the same brain area. These were checked against the raw scans and removed if they did not appear as “cells” during a qualitative evaluation.

### Outlying values

The selection of parameters for cell identification, the removal of the illumination artefact, the managing of areas with small volumes, and the removal of mis-labelled spots are procedures to limit as much as possible the presence of outliers. Despite these efforts, residual biological / technical outliers could be expected, either at the single cell level (e.g. unspecific antibody binding) or at the brain area level (e.g. disproportionate activation). Due to the limited sample size and the batch effects, the identification of outlying values was not trivial. We therefore chose to not use any rule (e.g. 3SD away from the mean) or statistical test to detect / exclude / replace outliers. Rather, we assumed that they may occur uniformly across samples, thereby giving rise to increased variation. To mitigate their effects, we used medians and quantiles rather than means to summarize the data.

### Missing values

The main source of missing value was the loss of animals due to insufficient staining quality (see ‘Experimental Design’ in Methods).

A second source of missing values was due to damaged brain areas during the experimental procedure. c-fos+ cells were counted per brain area across the whole brain. Damaged areas were manually detected in the 488nm image stack independently by at least two of three researchers (VB, HS, RD). The researchers were blinded to the experimental condition, and discrepancies were resolved with discussion. c-fos+ cells of damaged areas were removed, and then imputed by mirroring the xyz cells’ coordinates of the same brain area of the opposite hemisphere. Although this approach inherently assumes no differences between hemispheres, we preferred it to a multiple imputation approach because it did not require batch effects’ mitigation and it could be performed at a single cell rather than at brain area level.

A third source of missing values was linked to cell detection. The cell detection algorithm requires the definition of a minimum c-fos+ intensity. This parameter was optimized in pilot experiments, and it was kept identical throughout all experimental brains. In principle this is not a problem, since by rigorous standardization it is possible to mitigate batch effects and obtain comparable relative statistics. However, when a brain area has no active cells at t_0_, it needs to be further evaluated to conduct a proper standardization. In our experiment, two brain areas (FRP and AHN) had no c-fos+ cells in one t_0_ sample. Since this occurred only in one sample, we considered these brain areas as missing, not as zeros for analyses that required standardization with ratios to baseline.

### Preprocessing for region-based statistics

Additional pre-processing is required for region-based analyses. In Figure 1, we summarize which pre-processing steps were required for which type of analysis.

In region-based analyses, the total number of c-fos+ cells (i.e. absolute counts) was calculated per brain area. However, cell counts are by definition not normally distributed; rather, they follow a Poisson or (negative) binomial distribution. We therefore applied a Box-Cox transformation per block (i.e. each set for four different time-points), so that our data would resemble a normal distribution.

Different brain areas have different sizes; therefore, absolute counts of c-fos+ cells are not indicative of how active a certain brain area is. In analysis where different brain areas are compared, absolute counts need to be normalized to the size of the brain area. We therefore calculated the number of c-fos+ cells per thousand of the total cells in each brain area, by adapting the atlas by Erö and colleagues ^31^. We used the total cell count estimation rather than that of only neurons because several publications have reported c-fos+ glia and astrocytes (for a review, see ^32^), and it is in agreement with the presence of c-fos+ cells in the fiber tracts of our own data.

The number of c-fos+ cells differed across batches, although the relationship across time points was consistent within batches. Therefore, a normalization step was required to make the data more comparable across batches. Z-score normalization was performed per block, i.e. a unit of one sample per experimental group. With z-score normalization, the data is scaled with a mean of 0 and a standard deviation of 1, according to the formula (*x* - μ)/*σ*, where *x* is the observed value, μ is the mean of the sample, and *σ* corresponds to the standard deviation of the sample.

### Region-based analyses: active brain regions

With the exception of the single-cell strategy analysis, the analyses were planned on a subset of data (n=3), and later extended to the full dataset. The experimenter coding the analyses (VB) was blinded to the experimental groups.

We tested which brain areas had a significant increase from baseline in c-fos+ cell count per thousand of total cells (n_cfos+ / tot_). The dependent variable was scaled and normalized as explained in the data pre-processing section. We performed pairwise comparisons (Welch t-test, one-sided, alpha = 0.05) for each time point (t_30_, t_90_, t_180_) against t_0_. P-values were adjusted with the Benjamini-Hochberg (BH) procedure. For visualization only, we transformed the p-values with a -log_10_ transformation, and we grouped the brain areas according to the ABA embryological origin.

Next, we tested which brain areas were most active. The analysis was independently performed per block; therefore, no other batch-effect correction step was taken. For each block, we calculated the top 5% of n_cfos+ / tot_ independently of time point, and thereby identified per block the most active brain areas. Next, we counted how often a specific brain area was categorized as most active. We considered a brain area to be consistent across samples if it was present in at least 5 out of 9 blocks in a particular time point. If we consider the process to be random under the null hypothesis, the probability of a brain area to be present in 5 out of 9 blocks would be 0.1%. This probability was estimated with a simulation study. We simulated 1000 independent experiments, and each experiment consisted of 4 time points with 9 independent iterations (i.e., n_blocks_ = 9). For each iteration, we selected 18 brain areas, meaning the 5% of 90 (nbrain areas) * 4 (n_time points_). Each brain area had an equal probability of being selected (i.e., uniform distribution), and a brain area could be picked multiple times within each iteration (i.e., block), up to 4 (n_time point_ = 4). Then, we calculated across 1000 experiments the probability of a brain area to be in the top 5% of the distribution. This simulation gave information about how likely it is that the representation in the top 5% was chance.

Since c-fos is not uniformly distributed across the brain, we performed a simulation study to assess whether the pattern obtained was due to the baseline spatial distribution of c-fos. We downloaded via the ABA’s API the mRNA c-fos expression levels of 3 experiments that passed the ABA quality check (id: 80342219; 79912554; 68442895). We calculated the mean and standard deviation for each of the brain areas available (n_brain areas_ = 8). Since the resolution available for c-fos expression is lower than the resolution in our dataset, we assumed that the c-fos expression available corresponded to the location and scale parameters of a normal distribution defined by all the sub-areas. In other words, for each sub-area we sampled values from a normal distribution with location and scale parameters equal to those derived from the ABA atlas. This was performed for 9 independent samples (n_samples_ = 4) for each time point (n_time point_ = 4). Since the selection may not be linked only to baseline c-fos expression, but also to a natural increase due to the foot-shock, we multiplied the baseline expression levels with the overall increase in c-fos across time points in our experiment. For each brain area, we calculated the median of the ratio between each foot-shock time point (t_30_, t_90_, t_180_) and t_0_. This ratio was then multiplied by the estimated c-fos expression values. Of note, the ratio at t_0_ was always 1, meaning that the expression levels were estimated only from the ABA. We performed this simulation 1000 times, thereby simulating 1000 independent experiments. For each simulated experiment, we considered 9 blocks and 4 time points, as in our actual experiment. Then, for each block in each experiment, we selected the brain areas whose expression was in the top 5% of the distribution. Across the 1000 experiments, we then calculated the mean and standard deviation.

### Region-based analyses: order of activation

Since brain areas displayed a temporal dynamic pattern, we aimed to order the brain areas based on their c-fos+ expression. Ordering brain areas based on the time of their activation is not trivial, especially since in 3D microscopy time is discrete (n_time point_ = 4) rather than continuous (as, for example, in fMRI). Additionally, c-fos protein is not transient, but it peaks ~90 min and decays ~180 min after a stimulus. This dynamic may even be different depending on the brain area ^5^. With these challenges in mind, we aimed to analytically create a pseudo-time to increase the temporal resolution, which would in turn allow to order brain areas.

Among the approaches considered (Supplementary Table 3), we ultimately ordered areas based on the estimated time of mid-activation across blocks. c-fos+ cell counts were Z-transformed. Then, for each brain area we calculated the median across blocks of each time point. We interpolated a linear model between each two consecutive time-points: this line is the “continuum” of pseudo-time. To order the brain areas, we then considered at which pseudo-time point, c-fos activation reached its mid-activation level. Since the data was Z transformed, this corresponded to reaching the value 0. To limit the sensitivity of the pseudo-time, we binned the pseudo-time variable in bouts of 10 minutes. Each brain area was therefore grouped to the closest bout (binning). This approach has the advantage to create a criterion on which to categorize brain areas, but it does not consider the range (error) among which it could happen. The approach is ideal for areas that have one point of activation; it is biased for brain areas with a biphasic activation (e.g., at the beginning and at the end of our time curve).

For interpretation and visualization purposes, we classified the brain areas (Supplementary Table 4) according to functional networks relevant to stress. We followed Henckens and colleagues’ ^9^ results, to which the an amygdalar group was added.

### Combining voxel based and single cell analysis: sub-brain areas

We questioned whether c-fos+ cells are uniform within a brain area, or whether there are locations in which c-fos+ cells are most dense, i.e., are in closer proximity to each other. For this, we selected a brain area important for the stress response, i.e. the basolateral amygdala (BLA). All other brain areas can be visualized on the *abc4d* app.

For each sample separately, we estimated the probability density of the BLA at each cell, by using a kernel density estimation with Gaussian function. Kernel densities are routinely used to smooth data from a finite sample to make inferences about a population. Here, they were used to estimate the cell density within a brain area. Next, we filtered the cells with highest density per sample.

The BLA was divided into voxels of 30μm per side. Considering that there could be an alignment error of (maximum) ~23μm (Supplementary Figure 2b), we considered 30μm the minimum, interpretable size. To look for consistency across samples, we calculated how many samples had at least one cell in each voxel, and considered 3 the minimum for consistency. In each xyz direction, we calculated per time point the median and interquartile of the voxels’ position. Of note, due to the batch effects, calculating number of cells (or other measures of activation) across samples would have had little value, and would need to be standardized. We therefore opted for this more straightforward approach. To determine whether our observations were due to a chance process, we randomly attributed each sample to a time point, and perform the exact same analysis.

### Combining region-based & single cell analysis: Strategy

We hypothesized that brain areas can show activation with different strategies. With the “count strategy”, a brain area increases the *number* of c-fos+ cells with a low c-fos expression (intensity); with an “intensity strategy”, c-fos+ cells increase in *intensity* rather than number.

To test this hypothesis, we analyzed t_0_ samples, where we calculated the n_cfos+_ of each brain area, as well as the mean intensity of the cells in that area (intensity refers to the maximum intensity as reported by Imaris). Here, the mean rather than the median was intentionally used to be able to observe the increase in intensity due to a subgroup of cells within a brain area. For this analysis, we did not perform a batch effects correction; rather we took advantage of the differences across blocks. In our cell detection methodology (‘Image processing: cell detection’ in Methods), cells are identified as c-fos+ depending on intensity. This relationship should always be the same, irrespective of batch effects or time points. To quantify the relationship analytically, we therefore interpolated a linear model between the raw c-fos+ cell count and median intensity, of all brain areas of all samples.

The linear model was used as a discriminant criterion to classify whether a brain area had a strategy more towards intensity (above the regression line) or towards count (below the regression line). This categorization was performed for each brain area of each t_0_ sample (n_animal_ = 9) independently. We then calculated the probability of a brain area to belong to a certain categorization. The resulting variable was continuous between the values of 0 (i.e., all samples had an intensity strategy) and 1 (i.e., all samples had a count strategy).

If the categorization of brain areas would be a random process, the probability of brain areas to belong to a certain categorization would be normally distributed around μ = 0.5 under the null hypothesis (i.e. brain areas do not have a strategy). To validate that the null hypothesis would indeed follow a normal distribution, we performed a simulation study. In this study, we used the exact same analysis, but the values for intensity and c-fos+ cell count were drawn independently from a Poisson distribution *P*(*λ* = 3.5). The value for lambda *λ* was selected by qualitatively comparing the distribution of intensity and count in the data with computer generated Poisson distributions with different lambdas. However, the interpretation would not change if different values of lambda would be selected.

Next, we questioned whether brain areas may change strategy after stress, relatively to t_0_. Our experiment was not powered sufficiently to answer this particular research question, and therefore results should be interpreted as exploratory. From the categorization probability, we selected those brain areas that were consistent across samples, i.e. that had a specific categorization in at least 6 out of 9 samples. Of these, we selected those with a consistent change (increase or decrease) in count and / or intensity in at least one of the foot-shock time points (t_30_, t_90_, t_180_). We calculated the pairwise difference between t_0_ and each foot-shock time point for count as well as intensity. From this, we calculated the rate of change (count over intensity) per block for each time point. To compare data across brain areas, we converted the rates across samples to standardized mean differences (Hedge’s g). We then classified brain areas as having changed strategy after stress if the effect size was below 0.5 or higher than 2, meaning that the relative increase in activity must have doubled towards intensity or towards count after stress compared to baseline.

### Software

We developed the R package *abc4d* (“Analysis Brain Cellular activation in 4 Dimensions”) to ease the implementation of data pre-processing and analysis. Furthermore, we developed a web tool (https://vbonapersona.shinyapps.io/brain_after_footshock/) to interactively visualize the effects of acute stress on the brain area of choice.

All analyses were conducted with R (version 4.0.0) in the R studio environment on a macOS Mojave (version 10.14.6). The following R packages were core to this study: 1) *tidyverse* (version 1.3.0) for general data handling and visualization; *shiny* (v 1.6.0) for the generation of the web interface; *ComplexHeatmap* (v 2.4.3) for heatmap visualization.

## Data Availability

The data that supports the findings of the current study can be downloaded at https://osf.io/8muvw/. All figures can be reproduced from the data by following the guidelines provided alongside the code at https://osf.io/8muvw/.

## Code Availability

All code used in this manuscript is available at https://osf.io/8muvw/.

