## Supplementary information for "The mouse brain after foot-shock in 4D: temporal dynamics at a single-cell resolution"

This PDF contains:

Supplementary Note 1

Supplementary Figures 1-6

Supplementary Tables 1-4

The R package is available at:

The data is available at:

The interactive visualization is available at:

### Supplementary Notes

**Supplementary Note 1: c-fos**

c-fos is a proto-oncogene of the Fos family, which acts as a transcription factor upon heterodimerization with a member of the Jun family (Hughes & Dragunow, 1995; Kouzarides & Ziff, 1988). With the exception of a few constitutively active brain areas, c-fos is not expressed under baseline, i.e. non-stressed, circumstances (Luckman et al., 1994) but transiently induced by mild-to-severe acute stimuli, with activity-dependent intensity (Krisztina J Kovács, 1998).

### Supplementary Figures

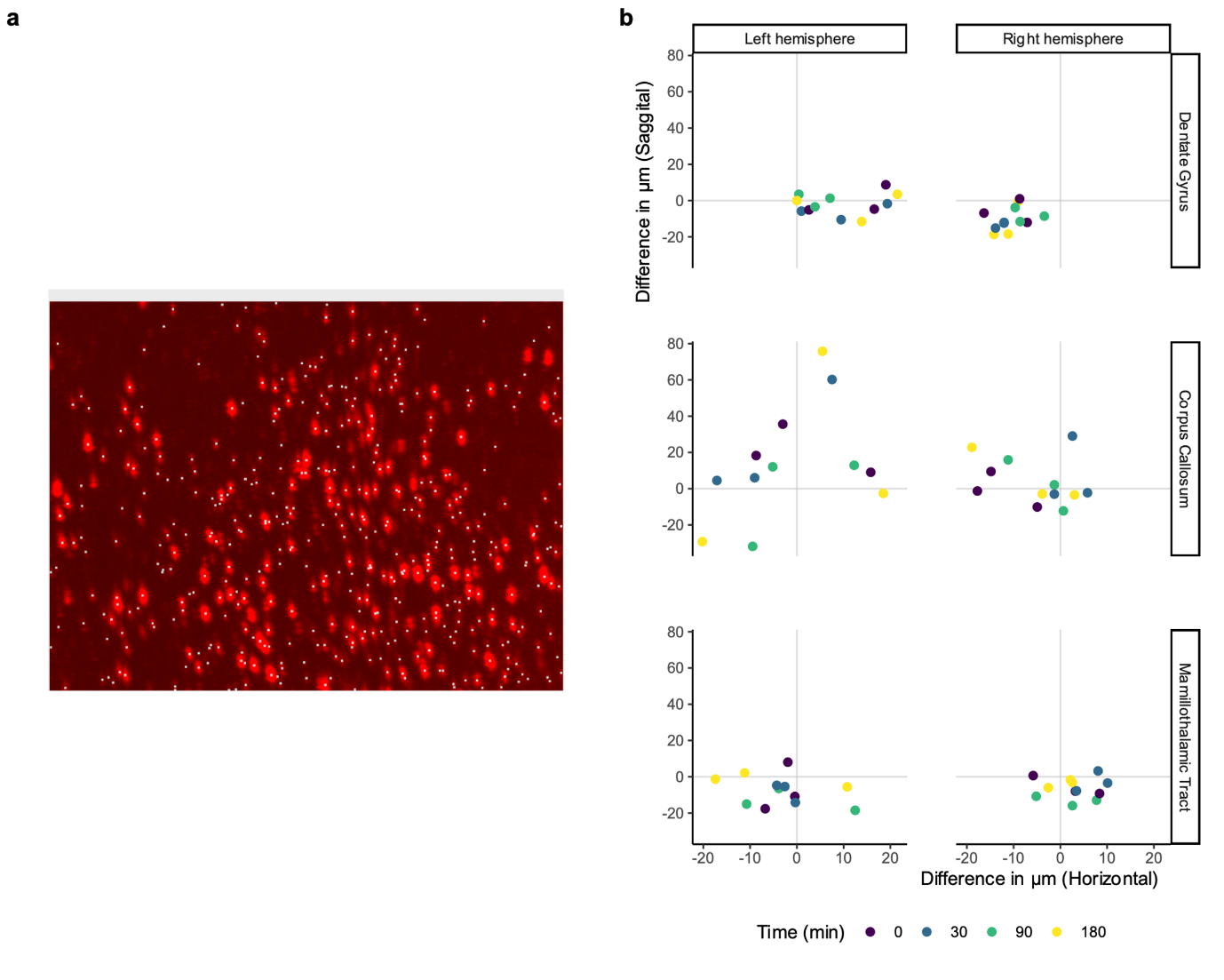

**Supplementary Figure 1. Cell detection and alignment validation.** a) Representative example of staining of c-fos+ cells (bright red). White squares represent objects identified by the Imaris algorithm as cells. b) Validation of alignment. Error of the alignment represented as distance between real and aligned objects along the horizontal and sagittal axis. Error was calculated in three separate brain areas (horizontal facets) for n = 3 samples per time point.

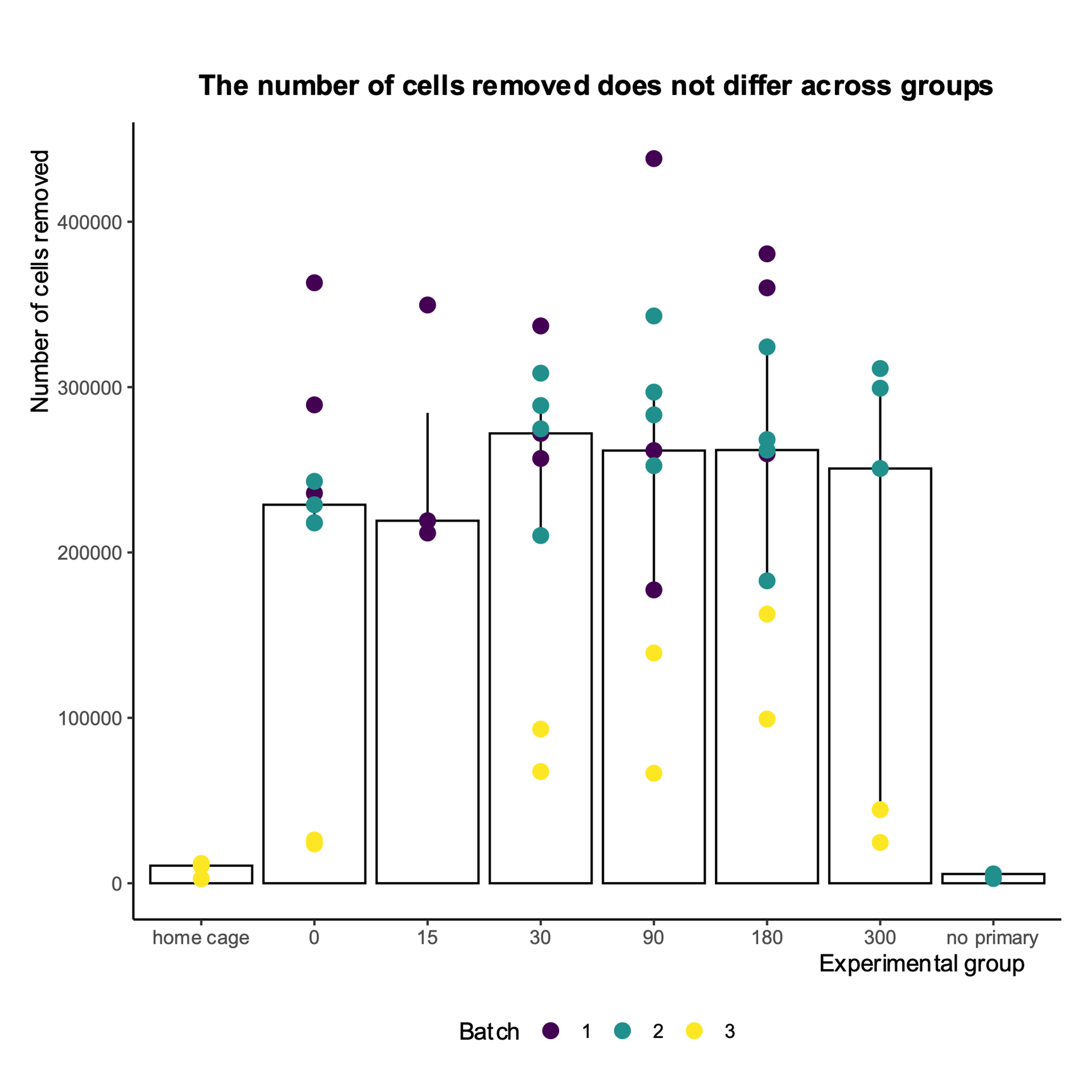

**Supplementary Figure 2. Data cleaning.** Number of ‘cells’ removed during the data cleaning procedure across all groups. Each dot corresponds to one sample. Data presented as median and IQR. Of note, t_15_ and t_300_ were only investigated in control experiments, and not across all batches (Supplementary Figure 3). Sample sizes (n): n_home cage_ = 3; n_0_ = 9; n_15_ = 3; n_30_ = 0; n_90_ = 9; n_180_ = 9; n_300_ = 6; n_no primary_ = 3.

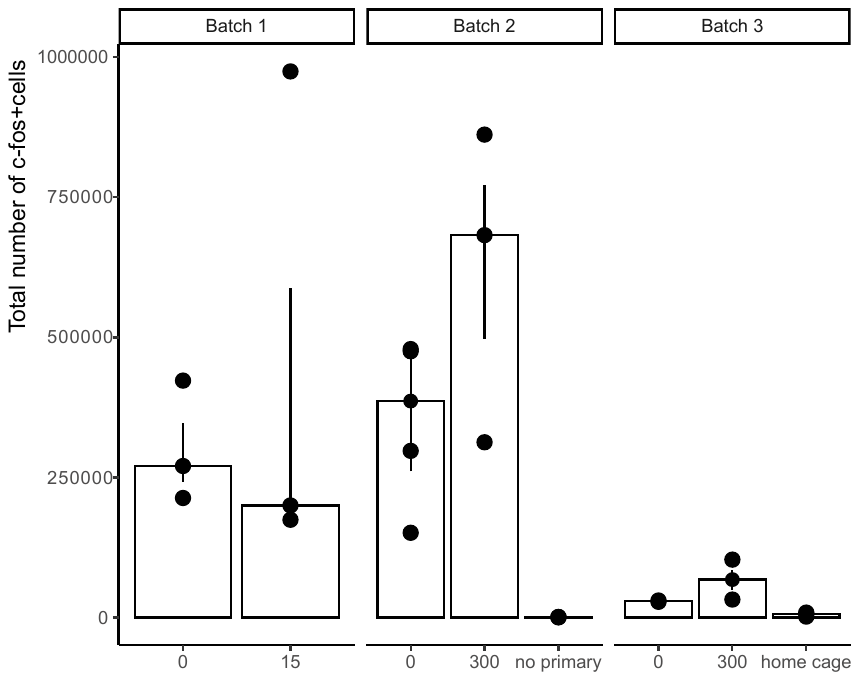

**Supplementary Figure 3. Total number of c-fos+ cells of control experiments.** 15 min after foot-shock is insufficient to detect an increase in c-fos expression. At 300 minutes, c-fos+ cell count is comparable to t_0_. Home cage group has lower c-fos+ cell count than the respective t_0_ group. No primary antibody group has nearly no counts. Each dot represents a sample, with the bar indicating the median, and the errorbar the interquartiles (IQR). The control experiments are represented separately with t_0_ groups of the same batch. Sample sizes (n): n_0_ = 9 (3 + 4 + 2 in each batch); n_15_ = 3; n_300_ = 6; n_no primary_ = 3; n_home cage_ = 3.

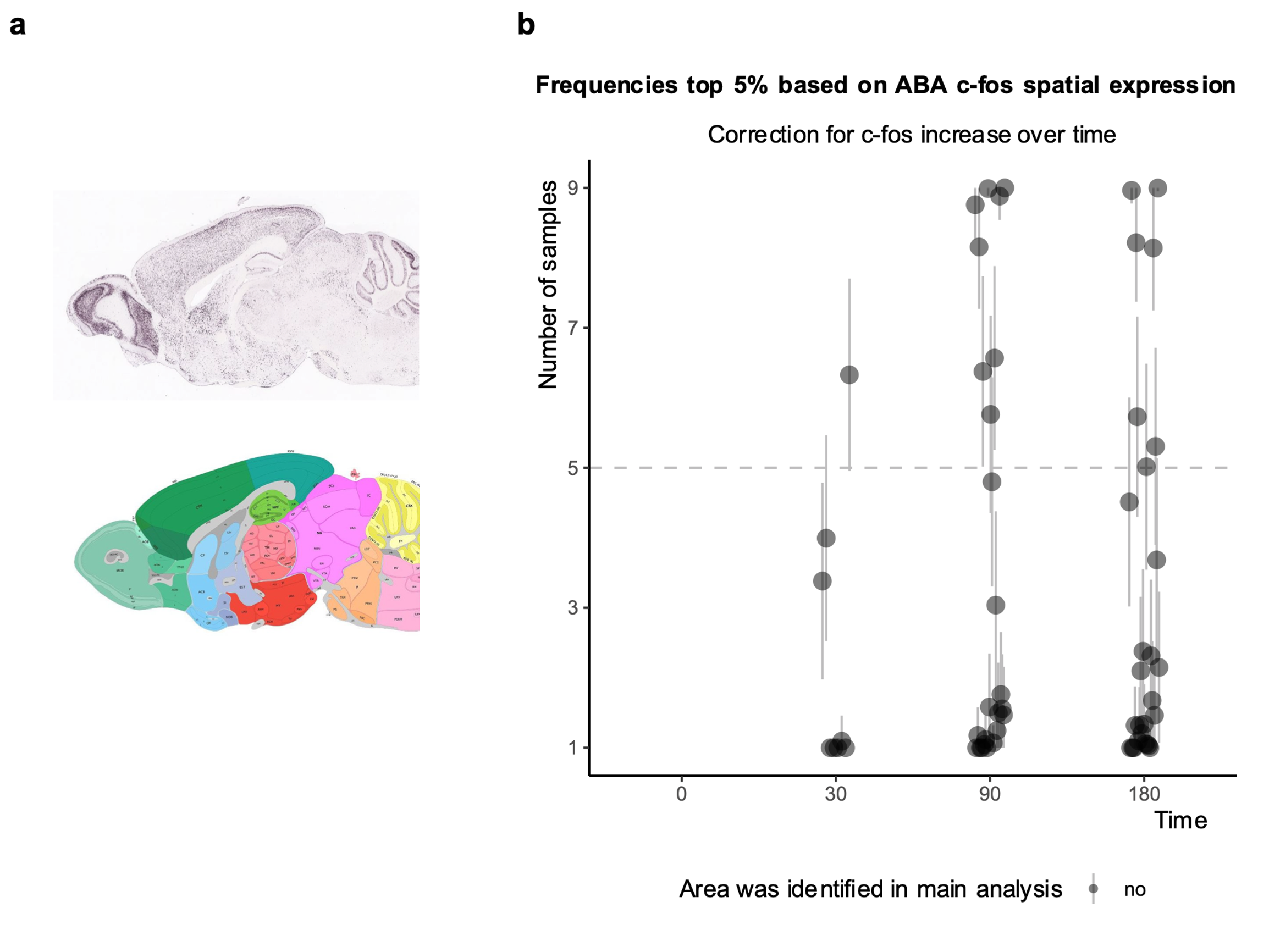

**Supplementary Figure 4. Simulation of most active brain areas based on c-fos ABA mRNA expression and increase of c-fos+ cells over time.** a) Regional distribution of c-fos as displayed by the Allen Brain Atlas (ABA). Red: hypothalamic areas. b) Results of the simulation study. An *in silico* dataset was created by using a sampling approach. c-fos mRNA expression values were downloaded from all the experiments available at the ABA API (n_experiment_ = 3). These values were used to sample weights to mimic what one would expect if the data were only due to c-fos expression and increase of c-fos over time. The procedure was repeated 1000 times. Each dot corresponds to a brain area that was present in the top 5% of the c-fos+ cell counts (per thousand of total) in one block. The vertical line of each dot represents the 95% confidence interval across the 1000 simulated experiments. None of the brain areas that were consistent in more than 5 samples was present in the actual experimental data.

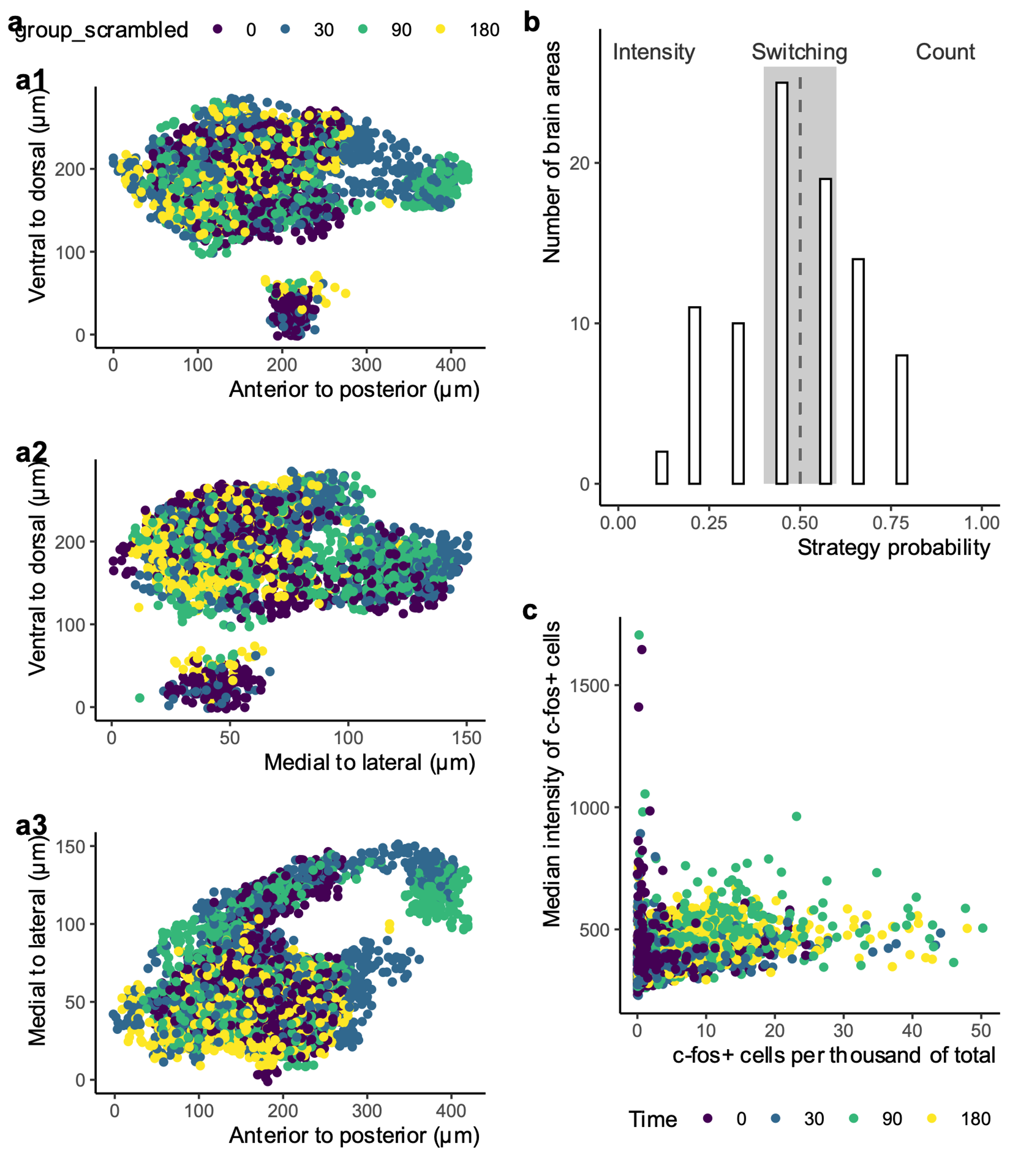

**Supplementary Figure 5**. a) c-fos+ cells in highly dense areas across scrambled time point. a1, a2, a3 refer to the different 2D views of the xyz coordinates. Time points were randomly allocated for each sample, so that each block (i.e. a unit of 4 time points) had still one sample per time point. b) Strategy probability of brain areas according to our hypothesis, i.e. the relationship between count and intensity is due to the technical set-up. c) Relationship between c-fos+ count and median intensity. Contrary to expectations, count of c-fos+ cells and mean intensity are not correlated to each other.

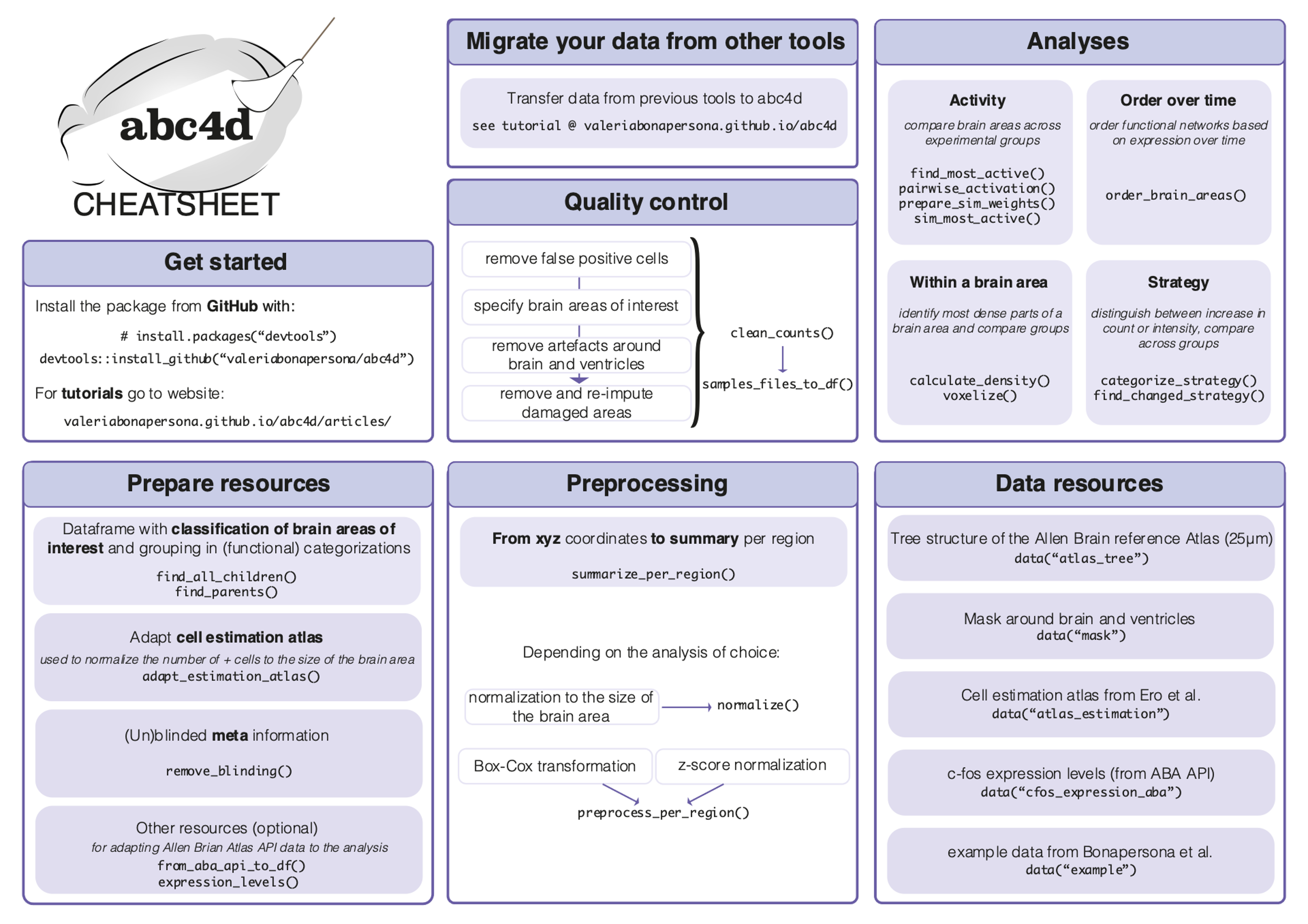

**Supplementary Figure 6**. Cheatsheet of *abc4d* package.

### Supplementary Tables

**Supplementary Table 1. Missing values.** a) List of missing animals with reasons. b) Damaged brain areas. These were removed from the analysis and re-imputed.

a) Missing animals

| **Batch** | **Animals missing** | **Reason** |
| --- | --- | --- |
| 2 | 11 male animals, 40 female animals | Staining faded. |
| 3 | 3 male animals | Staining faded (2 animals), scanning mistake (1 animal).  The 3 animals belonged to two separate blocks. To not exclude both blocks completely, we merged the remaining sample into one block by selecting the best quality stainings. |
| **Missing brain areas** |  |  |
| Sample | Damaged brain areas |  |
| 106 | SI right, FS right, SUBv-sp right |  |
| 107 | ACB left, OT left, PIR left, CP left, SI left |  |

b) Damaged areas

| **Sample ID** | **Damaged areas** |
| --- | --- |
| 13 | AAA left, CP left, OT right, AAA right, CP right, ENTl right |
| 14 | CP left, AAA left, VISC right, AIp right |
| 15 | MOp right, SSp right |
| 16 | PIR left, ENTl left, PL right, MOs right |
| 18 | CTX left |
| 19 | MOp left, AId left, GU left, AIv left |
| 21 | SSs left, CP left, PAR left, HPF left, AAA right |
| 22 | RSPd left, VISp left, PRE left, AAA right |
| 23 | RSPd left, RSPagl left, VISpm left, RSPv left |
| 24 | SI left, FS left, CP left, AAA right, CP right |
| 25 | PTLp left |
| 26 | PAG left, ICe left, SCs left |
| 27 | SSs left, PAA left, PIR left, COApl left, COAa left |
| 34 | AAA right, CP right, SSs right, MEAav right, PERI right, ECT right |
| 35 | AAA right, CP right, CEAm right, AUDd right, VISl right, TEa right, PRE right, ec right, dhc right, PRE right, SUBv right |
| 36 | PERI left , AIp right, CP right, AAA right |
| 38 | PERI left , ENTl left |
| 39 | VISpl left , VISp left , ec left , dhc left , PRE left , MOs , MOp , TEa , ECT , ENTm , PAR , ec , dhc |
| 40 | VISC left, AIp left, CP left, ECT right, ec right, dhc right, PAR right, ENTl right |
| 41 | OT left, PIR left, CP left, OT right, PIR right, AAA right, FS right, CP right, AIp right |
| 42 | MOs left , RSPv left , RSPd left , NA left , MOs right, TEa right, ECT right, PERI right, SUBv-sp right, ENT right, PAR right |
| 43 | OT left , SI left , FS left , CP left , PTLp right, TEa right |
| 44 | AId left, MOp left, ORBl left, CP left, OT left, FS left, CP left, VISpl left, ec left, dhc left, POST left, OT right, FS right, CP right |
| 45 | OT left , FS left , CP left , ECT left , OT right, FS right, CP right |
| 46 | OT left, FS left, CP left, CP right, OT right, FS right, AUDd right, AUDpo right, ECT right |
| 47 | TTd left , AON left , AAA left , CP left , IA left , ENTl left , ECT left , CP right, AAA right, IA right, ENTl right, ECT right |
| 48 | OT left, SI left, ACB left, CP left, AIp left, OT right, SI right, FS right, CP right, ECT right, ENTl right |
| 49 | VISpl left, ECT left, RSPd right, RSPv right |
| 50 | AId left, AON left, PIR left, AIv left, GU left, AAA right, CP right |
| 51 | AAA left, CP left, VISp left, VISal left, VISam left, VISp left, VISl left, POST left, PRE left, AAA right, CP right, AUDd right, PTLp right, ENT right, PAR right |
| 51 | SSs left, CP left, ECT left, TEa left, ENTm right |
| 106 | SI right, FS right, SUBv-sp right |
| 107 | ACB left, OT left, PIR left, CP left, SI left |
| 108 | SSp left , GU left , OT right, SI right, ACB right, CP right, POST right, ec right, VISpm right, VISp right, RSPv right, RSPd right |
| 109 | MOs right |
| 112 | ECT right, PERI right, ENTl right |
| 113 | VISpm left, VISp left, ec left, dhc left, POST left, MOs right, MOp right, PRE right, HPF right, SUBv-sp right, alv right, ec right |
| 115 | OT left |
| 116 | SUBv-sp right |
| 117 | PIR right, PAR right, ENTm right, ENTl right |
| 118 | SSs left , FS left , act left , CP left |
| 119 | SSp right |
| 120 | CP left |
| 121 | CP right |
| 122 | TEa right, SUBv-sp right, PAR right |

**Supplementary Table 2. List of brain areas included in the analysis.** The categorization follows the structure and acronym of the Allen Brain Reference atlas (25µm). ID is the code used by the ABA.

| **ID** | **Name brain area** | **Acronym** |
| --- | --- | --- |
| 23 | Anterior amygdalar area | AAA |
| 31 | Anterior cingulate area | ACA |
| 56 | Nucleus accumbens | ACB |
| 88 | Anterior hypothalamic nucleus | AHN |
| 95 | Agranular insular area | AI |
| 223 | Arcuate hypothalamic nucleus | ARH |
| 239 | Anterior group of the dorsal thalamus | ATN |
| 247 | Auditory areas | AUD |
| 295 | Basolateral amygdalar nucleus | BLA |
| 319 | Basomedial amygdalar nucleus | BMA |
| 351 | Bed nuclei of the stria terminalis | BST |
| 382 | Field CA1 | CA1 |
| 423 | Field CA2 | CA2 |
| 463 | Field CA3 | CA3 |
| 776 | corpus callosum | cc |
| 536 | Central amygdalar nucleus | CEA |
| 583 | Claustrum | CLA |
| 631 | Cortical amygdalar area | COA |
| 672 | Caudoputamen | CP |
| 784 | corticospinal tract | cst |
| 726 | Dentate gyrus | DG |
| 830 | Dorsomedial nucleus of the hypothalamus | DMH |
| 856 | Thalamus polymodal association cortex related | DORpm |
| 864 | Thalamus sensory-motor cortex related | DORsm |
| 814 | Dorsal peduncular area | DP |
| 895 | Ectorhinal area | ECT |
| 909 | Entorhinal area | ENT |
| 942 | Endopiriform nucleus | EP |
| 958 | Epithalamus | EPI |
| 1000 | extrapyramidal fiber systems | eps |
| 184 | Frontal pole cerebral cortex | FRP |
| 998 | Fundus of striatum | FS |
| 1057 | Gustatory areas | GU |
| 1105 | Intercalated amygdalar nucleus | IA |
| 44 | Infralimbic area | ILA |
| 51 | Intralaminar nuclei of the dorsal thalamus | ILM |
| 59 | Intermediodorsal nucleus of the thalamus | IMD |
| 131 | Lateral amygdalar nucleus | LA |
| 138 | Lateral group of the dorsal thalamus | LAT |
| 896 | thalamus related | lfbst |
| 194 | Lateral hypothalamic area | LHA |
| 226 | Lateral preoptic area | LPO |
| 275 | Lateral septal complex | LSX |
| 290 | Hypothalamic lateral zone | LZ |
| 290 | Hypothalamic lateral zone | LZ |
| 323 | Midbrain motor related | MBmot |
| 331 | Mammillary body | MBO |
| 339 | Midbrain sensory related | MBsen |
| 348 | Midbrain behavioral state related | MBsta |
| 362 | Mediodorsal nucleus of thalamus | MD |
| 403 | Medial amygdalar nucleus | MEA |
| 991 | medial forebrain bundle system | mfbs |
| 500 | Somatomotor areas | MO |
| 515 | Medial preoptic nucleus | MPN |
| 904 | Medial septal complex | MSC |
| 619 | Nucleus of the lateral olfactory tract | NLOT |
| 698 | Olfactory areas | OLF |
| 714 | Orbital area | ORB |
| 754 | Olfactory tubercle | OT |
| 780 | Posterior amygdalar nucleus | PA |
| 788 | Piriform-amygdalar area | PAA |
| 818 | Pallidum dorsal region | PALd |
| 826 | Pallidum medial region | PALm |
| 835 | Pallidum ventral region | PALv |
| 843 | Parasubiculum | PAR |
| 922 | Perirhinal area | PERI |
| 946 | Posterior hypothalamic nucleus | PH |
| 972 | Prelimbic area | PL |
| 1037 | Postsubiculum | POST |
| 1084 | Presubiculum | PRE |
| 1109 | Parastrial nucleus | PS |
| 63 | Paraventricular hypothalamic nucleus descending division | PVHd |
| 141 | Periventricular region | PVR |
| 149 | Paraventricular nucleus of the thalamus | PVT |
| 157 | Periventricular zone | PVZ |
| 165 | Midbrain raphe nuclei | RAmb |
| 254 | Retrosplenial area | RSP |
| 262 | Reticular nucleus of the thalamus | RT |
| 453 | Somatosensory areas | SS |
| 502 | Subiculum | SUB |
| 541 | Temporal association areas | TEa |
| 877 | tectospinal pathway | tsp |
| 589 | Taenia tecta | TT |
| 614 | Tuberal nucleus | TU |
| 629 | Ventral anterior-lateral complex of the thalamus | VAL |
| 669 | Visual areas | VIS |
| 677 | Visceral area | VISC |
| 685 | Ventral medial nucleus of the thalamus | VM |
| 693 | Ventromedial hypothalamic nucleus | VMH |
| 709 | Ventral posterior complex of the thalamus | VP |

**Supplementary Table 3**. List of analytical approaches considered for ordering brain areas based on c-fos activation.

| **Approach** | **Brief explanation** | **Not pursued because:** |
| --- | --- | --- |
| Clustering | Clustering to reduce dimensions, then order the cluster groups. The pseudo-time would then have resolution equal to the number of clusters. Ordering could be achieved by comparing to a simulated model of possible clusters out of theory (example: cluster with only initial activation at t_30_; cluster with activation at t_90;_ clustering with activation at t_90_ as well as t_180_). | All brain areas were activated; therefore, very minimal difference would appear between clusters. Furthermore, creating “expected” cluster models is not trivial. |
| Derivatives | Identify the steepest derivative between each two consecutive time points. This can be performed per sample (probabilistic approach) or on the median across samples. It might be able to identify multiple activations (e.g. if first and third derivatives are steeper than the second). | Too many rules (e.g. only one derivative is the steepest, two derivatives are the steepest…), therefore it has the same problem as creating the “expected cluster” model as described above. Furthermore, very pseudo-time resolution (n = 3). |
| Peaks | Fit a *loess* curve for each sample and identify the maxima. Each maxima is considered a peak. Advantage that it can identify multiple activations for a single brain area | Since all brain areas were so activated, many peaks would appear within the same range (poor pseudo-time resolution). As a consequence, too much importance would be given to the type of curve used to fit the data. |

**Supplementary Table 4. Functional categorization of brain areas.** Brain areas were classified in functional groups relevant to the stress response, by adapting (Henckens et al., 2015). *cx = cortex.*

| **Functional categorization** | **Brain area** | **Acronym** |
| --- | --- | --- |
| Amygdala | Anterior amygdalar area | AAA |
| Amygdala | Basolateral amygdalar nucleus | BLA |
| Amygdala | Basomedial amygdalar nucleus | BMA |
| Amygdala | Central amygdalar nucleus | CEA |
| Amygdala | Cortical amygdalar area | COA |
| Amygdala | Intercalated amygdalar nucleus | IA |
| Amygdala | Lateral amygdalar nucleus | LA |
| Amygdala | Medial amygdalar nucleus | MEA |
| Amygdala | Posterior amygdalar nucleus | PA |
| Amygdala | Piriform-amygdalar area | PAA |
| Hippocampus | Field CA1 | CA1 |
| Hippocampus | Field CA2 | CA2 |
| Hippocampus | Field CA3 | CA3 |
| Hippocampus | Dentate gyrus | DG |
| Hippocampus | Entorhinal area | ENT |
| Hippocampus | Parasubiculum | PAR |
| Hippocampus | Postsubiculum | POST |
| Hippocampus | Presubiculum | PRE |
| Hippocampus | Subiculum | SUB |
| Hypothalamus | Anterior hypothalamic nucleus | AHN |
| Hypothalamus | Arcuate hypothalamic nucleus | ARH |
| Hypothalamus | Dorsomedial nucleus of the hypothalamus | DMH |
| Hypothalamus | Lateral hypothalamic area | LHA |
| Hypothalamus | Lateral preoptic area | LPO |
| Hypothalamus | Hypothalamic lateral zone | LZ |
| Hypothalamus | Hypothalamic lateral zone | LZ |
| Hypothalamus | Mammillary body | MBO |
| Hypothalamus | Medial preoptic nucleus | MPN |
| Hypothalamus | Posterior hypothalamic nucleus | PH |
| Hypothalamus | Parastrial nucleus | PS |
| Hypothalamus | Paraventricular hypothalamic nucleus descending division | PVHd |
| Hypothalamus | Periventricular region | PVR |
| Hypothalamus | Periventricular zone | PVZ |
| Hypothalamus | Tuberal nucleus | TU |
| Hypothalamus | Ventromedial hypothalamic nucleus | VMH |
| Motor cx | Somatomotor areas | MO |
| Prefrontal cx | Anterior cingulate area | ACA |
| Prefrontal cx | Orbital area | ORB |
| Prefrontal cx | Prelimbic area | PL |
| Prefrontal cx | Taenia tecta | TT |
| Primary somatosensory cx | Somatosensory areas | SS |
| Thalamus | Anterior group of the dorsal thalamus | ATN |
| Thalamus | Thalamus polymodal association cortex related | DORpm |
| Thalamus | Thalamus sensory-motor cortex related | DORsm |
| Thalamus | Epithalamus | EPI |
| Thalamus | Intralaminar nuclei of the dorsal thalamus | ILM |
| Thalamus | Intermediodorsal nucleus of the thalamus | IMD |
| Thalamus | Lateral group of the dorsal thalamus | LAT |
| Thalamus | Mediodorsal nucleus of thalamus | MD |
| Thalamus | Paraventricular nucleus of the thalamus | PVT |
| Thalamus | Reticular nucleus of the thalamus | RT |
| Thalamus | Ventral anterior-lateral complex of the thalamus | VAL |
| Thalamus | Ventral medial nucleus of the thalamus | VM |
| Thalamus | Ventral posterior complex of the thalamus | VP |
| Visual cx | Visual areas | VIS |
